# Stable isotope labeling reveals preferential ammonium acquisition by hydroponic lettuce under ambient and elevated CO_2_

**DOI:** 10.64898/2026.11.16.767057

**Authors:** Luke L. Fountain, Sebastien Malbrough, Elison B. Blancaflor, Raymond M. Wheeler, Gioia D. Massa

## Abstract

Nitrogen (N) form acquired by plants reflects both physiological preference and environmentally mediated availability. Hydroponic controlled-environment agriculture (CEA) removes many constraints governing N availability yet remains predominantly reliant on nitrate (NO_3_^-^) nutrition. We used differential ^15^N stable-isotope labeling to quantify ammonium (NH_4_^+^) and NO_3_^-^ acquisition by four lettuce (*Lactuca sativa* L.) cultivars grown hydroponically under terrestrial ambient (410 ppm) and spaceflight-relevant elevated (3000 ppm) CO_2_. Plants received equivalent concentrations of NH_4_^+^ and NO_3_^-^, with either N form selectively enriched with ^15^N during the final 24 h of growth. All cultivars exhibited significantly greater shoot ^15^N enrichment following ^15^NH_4_^+^ than ^15^NO_3_^-^ labeling under both CO_2_ environments, demonstrating consistent preferential NH_4_^+^ acquisition. Preference magnitude varied among cultivars and broadly corresponded with biomass and total shoot N accumulation, driven predominantly by increased NH_4_^+^-derived enrichment rather than reduced NO_3_^-^-derived enrichment. Elevated CO_2_ did not significantly alter N-form preference, although strong NH_4_^+^ preference persisted at 3000 ppm. These findings distinguish physiological acquisition preference from environmentally mediated N availability and provide a basis for reconsidering nitrate-dominant nutrition in terrestrial and space CEA, including integration with recycled reduced N.

**Summary statement:** Differential ^15^N labeling revealed consistent preferential ammonium acquisition across four hydroponically-grown lettuce cultivars under both ambient and spaceflight-relevant elevated CO_2_, challenging the assumption that nitrate-dominant nutrition reflects lettuce N-form preference

## Introduction

Nitrogen (N) is an essential macronutrient required for plant growth and is acquired predominantly as the inorganic ions ammonium (NH_4_^+^) and nitrate (NO_3_^-^). Although plant species are frequently characterized as preferentially acquiring one N form over another, apparent N-form preference is strongly influenced by the environment in which N is supplied. In soils, the relative availability of NH_4_^+^ and NO_3_^-^ is governed by interacting biological and physicochemical processes, including mineralization, nitrification, denitrification, microbial competition, ion mobility and retention, which are themselves influenced by environmental conditions such as soil moisture, oxygen availability, temperature and pH (Britto & Kronzucker, 2013; Chalk & Smith, 2021). Consequently, the N form predominantly acquired by a plant under field conditions may reflect the N form made most accessible by its environment as much as an intrinsic physiological preference for that N source.

Evidence from natural ecosystems demonstrates the importance of this environmental control. Across a tropical forest precipitation gradient, Houlton et al. (2007) observed a community-wide shift from predominantly NO_3_^-^ acquisition in drier soils to NH_4_^+^ acquisition in wetter soils, closely corresponding with changes in the relative abundance of the two inorganic N pools. Similarly, Wang and Macko (2011) demonstrated that plant N-form preference varied across an aridity gradient and proposed that these differences were constrained by climatic conditions and associated differences in the relative availability of NH_4_^+^ and NO_3_^-^. These observations illustrate the difficulty of separating physiological N-form preference from environmentally mediated N availability in soil-based systems. Indeed, NO_3_^-^ is highly mobile in soil and is commonly the principal inorganic N source in well-aerated agricultural soils, whereas NH_4_^+^ can dominate under contrasting environmental conditions (Wang & Macko, 2011). Thus, the widespread association between crop production and NO_3_^-^ acquisition does not necessarily establish that NO_3_^-^ is intrinsically the preferred N form when NH_4_^+^ and NO_3_^-^ are equally available.

Controlled-environment agriculture (CEA), and hydroponic cultivation in particular, provides an opportunity to disentangle physiological N acquisition from many of these environmental constraints. In recirculating hydroponic systems, NH_4_^+^ and NO_3_^-^ can be supplied directly to roots at defined concentrations without the soil processes that ordinarily determine their relative availability. Nevertheless, modern hydroponic nutrient formulations remain predominantly NO_3_^-^-based, with NH_4_^+^ generally supplied as a comparatively small proportion of total N (Sonneveld & Voogt, 2009; Voogt & Bar-Yosef, 2019). This partly reflects the operational and physiological challenges associated with higher NH_4_^+^ supply, which can result in rapid rhizosphere acidification, nutrient imbalances, and growth inhibition (Britto & Kronzucker, 2013; Natarajan Sakthivel et al., 2023). The predominance of NO_3_^-^ in hydroponic practice may therefore reflect the challenges associated with managing NH_4_^+^ as well as assumptions inherited from soil-based crop production, rather than physiological acquisition preference alone. Determining which N form plants preferentially acquire when both are supplied at equivalent concentrations and are continuously available is consequently important for establishing whether nitrate-dominant nutrition optimally aligns with crop N-acquisition physiology in engineered production systems.

There is a physiological basis to hypothesize that plants may preferentially acquire NH_4_^+^ when external availability of the two N forms is equivalent. Following uptake, NO_3_^-^ must first be reduced to NO_3_^-^ and subsequently NH_4_^+^ before N can be incorporated into organic compounds through the glutamine synthetase/glutamate synthase (GS/GOGAT) pathway. NH_4_^+^, by contrast, can enter this assimilation pathway directly, avoiding the additional energetic and reductant requirements associated with NO_3_ ^-^ reduction (Bloom et al., 1992). Although NH_4_^+^ nutrition imposes other physiological costs, and excessive NH_4_^+^ concentrations can become detrimental (Britto & Kronzucker, 2013), these limitations do not necessarily preclude preferential acquisition of NH_4_^+^ when both forms are present at non-toxic concentrations. The physiological preference expressed under controlled N availability may therefore differ from the apparent preference inferred from N acquisition in soil.

Atmospheric CO_2_ concentration may further modify this relationship. Elevated CO_2_ has been reported to inhibit shoot NO ^-^ assimilation in C plants, and plant growth responses to CO enrichment can consequently depend upon the N form supplied (Bloom et al., 2002; Bloom et al., 2012). Environments with highly elevated CO_2_ may therefore alter the relative physiological advantages associated with acquisition of oxidized and reduced N. This interaction is particularly relevant to space crop production, where plants may be cultivated under atmospheric CO_2_ concentrations substantially greater than terrestrial ambient conditions. Concentrations of several thousand ppm have historically occurred aboard the International Space Station (ISS), although CO_2_ concentrations vary spatially and temporally within the vehicle (Beard, 2020; Georgescu et al., 2020). Elevated atmospheric CO□ concentrations are likely to remain relevant to future space crop production systems, owing to the challenges of managing crew-generated CO within closed habitats and the potential use of elevated CO□ to enhance crop productivity (Maiwald et al., 2020; Ewert et al., 2022). Determining whether elevated CO_2_ alters physiological N-form preference is therefore necessary to establish whether N-acquisition strategies characterized under terrestrial conditions remain applicable to crops grown in spaceflight-relevant environments.

Nitrogen form is also important at the crop-production system level. Space-based CEA (SpaCEA) must provide sufficient nutrients to sustain crop productivity while minimizing resource requirements. The integration of crop production with waste and nutrient recovery has consequently been identified as an important step toward bioregenerative life-support systems (BLSS), with a recently proposed BLSS Readiness Level framework identifying nutrient-loop closure and subsequent recycled fertigation as key stages in developing crops as operational components of BLSS (Fountain et al., 2026). Human waste streams represent a potentially important source of recoverable N. Urinary N is predominantly excreted as urea, which can be hydrolyzed to NH_4_^+^, whereas production of NO_3_^-^ from this recovered reduced N requires subsequent oxidation through nitrification (Verbeelen et al., 2021). If candidate crops can utilize a substantial proportion of their N directly as NH_4_^+^, greater integration of crop production with waste-N recovery could potentially reduce the extent of nitrification required and associated energy and infrastructure requirements before crop delivery and decrease reliance on N fertilizer transported from Earth. Conversely, if plants intrinsically favor NO_3_^-^, even when NH_4_^+^ is similarly available, conversion of recovered N to NO_3_^-^ may represent a necessary component of regenerative nutrient management. Establishing physiological N-form preference is therefore important for rationally designing N-recycling strategies rather than assuming that nutrient forms conventionally used in terrestrial agriculture are necessarily optimal for SpaCEA.

The same question is increasingly relevant to terrestrial CEA. Hydroponic production has already decoupled crop nutrition from many of the soil processes that are primary determinants of inorganic N availability, yet NO_3_^-^-dominant nutrient strategies have largely persisted. At the same time, waste-derived nutrient streams such as anaerobic digestate, represent potential sources of NH_4_^+^-rich N that could be integrated with hydroponic crop production (van Rooyen & Nicol, 2022). Understanding whether crops preferentially acquire NH_4_^+^ when environmental constraints on its availability are removed could therefore inform both optimization of conventional hydroponic nutrition and development of more circular CEA systems capable of coupling nutrient recovery directly with crop production. Engineered cultivation environments provide an opportunity not simply to reproduce nutrient regimes inherited from soil-based agriculture, but to design nutrient supply around plant physiology and the resources locally available to the production system.

Here, we used differential ^15^N stable-isotope labeling to quantify the relative acquisition of NH_4_^+^ and NO_3_^-^ by four lettuce (*Lactuca sativa* L.) cultivars grown in recirculating hydroponic systems. Plants were supplied with NH_4_^+^ and NO_3_^-^ at equivalent concentrations while either the NH_4_^+^ or NO_3_^-^ pool was selectively enriched with ^15^N, allowing N-form acquisition to be compared under equivalent external supply. We further examined whether N-form preference was modified under an elevated CO_2_ concentration of 3000 ppm, selected as a spaceflight-relevant elevated concentration within the range historically encountered aboard the ISS (Beard, 2020), relative to terrestrial ambient CO_2_ (410 ppm). We hypothesized that lettuce would preferentially acquire NH_4_^+^ when both N forms were equally supplied and that this preference would be enhanced under elevated CO_2_ because of the additional reductant requirements associated with NO_3_^-^ assimilation. By separating N-form acquisition from many of the soil processes that ordinarily regulate N availability, this study aimed to determine whether nitrate-dominant nutrition reflects the intrinsic N-acquisition physiology of lettuce and to establish a physiological basis for reconsidering N management in terrestrial and SpaCEA.

## Materials and Methods

### Experimental design

The experiment was organized in a factorial design comprising four lettuce cultivars, two CO_2_ concentrations, and two ^15^N isotopic labeling treatments, resulting in 16 treatment combinations. Eight plants were established per treatment combination in each experimental run, with five plants targeted for individual isotopic analysis and the remaining plants maintained to accommodate potential germination or establishment losses. The experiment was conducted in three independent runs over time, with a target of 15 individual plants analyzed per treatment combination across the three experimental runs. Where plant loss resulted in fewer than five suitable biological replicates for a treatment combination in an earlier run, additional plants from the corresponding treatment combination were analyzed in the final run to achieve the target sample size.

Plants were maintained at their respective CO_2_ concentrations throughout the experimental period, whereas differential ^15^N stable isotope labeling treatments were imposed only during the final 24 h prior to harvest. At harvest, plant shoot biomass was recorded and shoot tissue was collected from each analyzed plant for determination of total N and ^15^N abundance by isotope-ratio mass spectrometry (IRMS).

### Growth conditions and plant material

Experiments were conducted in Percival (Perry, Iowa, USA) walk-in controlled-environment growth chambers under either 410 or 3000 ppm CO□. The 410 ppm treatment was maintained in a growth chamber equipped with active CO_2_ scrubbing, which was required because background CO_2_ concentrations within the facility frequently exceeded terrestrial ambient concentrations. The 3000 ppm treatment was maintained in a separate growth chamber. Consequently, CO_2_ treatments remained assigned to the same respective chambers across experimental runs, and growth chamber and CO_2_ concentration were confounded. Effects associated with CO_2_ concentration were therefore interpreted with appropriate caution. Chambers were maintained at a constant temperature of 23 °C and 70% relative humidity. Plants were illuminated using OSRAM Phytofy® LED fixtures at an average photosynthetic photon flux density (PPFD) of 300 µmol m ² s ¹ with a spectral composition of 49% red, 17% green, and 34% blue light and a 16-h photoperiod.

Four lettuce (*Lactuca sativa* L.) cultivars were used in this study: ‘Dragoon’ (Johnny’s Selected Seeds, Winslow, ME, USA), ‘Outredgeous’ (Johnny’s Selected Seeds), ‘Waldmann’s Dark Green’ (Johnny’s Selected Seeds), and ‘Black Seeded Simpson’ (Burpee, Warminster, PA, USA). Three seeds were sown directly into 2.5 × 2.5 × 5 cm horticultural foam cubes (Sublime®, BVB Substrates, the Netherlands) positioned within the hydroponic growth system and remained in the system from sowing until harvest at 22 days after planting (DAP). A humidity dome was maintained over emerging seedlings for the first 10 DAP.

Plants were grown in custom two-tier nutrient film technique (NFT) hydroponic systems. Each tier comprised four 5-ft (1.5 m) NFT channels (CropKing Inc., Ohio, USA), with eight plant positions per channel and a total capacity of 32 plants per tier. Each tier contained eight plants of each lettuce cultivar, with individual plants assigned to positions in a randomized layout within each tier for each experimental run. Nutrient solution was supplied from 20-L opaque reservoirs and pumped to the NFT channels using a submersible pump at an average flow rate of approximately 450 mL min ¹. Channels were mounted on transparent acrylic supports of progressively differing heights to provide a downward slope, with nutrient solution draining from the channels through 2.5-inch (6.35 cm)-diameter PVC piping and returning to the reservoir. Nutrient solution was continuously recirculated throughout the experimental period.

Plants were supplied with 1× Yoshida nutrient solution, prepared as described by Yoshida *et al*. (1976), which was selected because nitrogen is supplied exclusively as NH NO and therefore provides NH_4_^+^ and NO_3_^-^ at an equimolar ratio. The composition of the nutrient solution is provided in Table 1. Reservoirs were maintained at a working volume of 20 L throughout plant growth prior to isotopic labeling. Nutrient solution was maintained without replacement for the first 10 DAP and was completely replaced every 2 d thereafter to minimize changes in the relative availability of NH_4_^+^ and NO_3_^-^ resulting from plant nutrient uptake. On days when complete solution replacement was not performed, reservoir volume was restored to 20 L with fresh 1× Yoshida nutrient solution as required; supplementation was not required during the first 10 DAP. Nutrient solution pH was initially adjusted to 5.8 and subsequently readjusted to 5.8 daily using 1 M KOH to compensate for progressive solution acidification.

**Table 1:** Yoshida nutrient solution composition, adapted from Yoshida *et al*. (1976).

| **Element** | **Reagent** | **Final concentration of element in solution (ppm)** |
|---|---|---|
| **Macronutrients** |  |  |
| N | NH<sub>4</sub>NO<sub>3</sub> | 40 |
| P | NaH<sub>2</sub>PO<sub>4</sub>·2H<sub>2</sub>O | 10 |
| K | K<sub>2</sub>SO<sub>4</sub> | 40 |
| Ca | CaCl<sub>2</sub>·2H<sub>2</sub>O | 40 |
| Mg | MgSO<sub>4</sub> | 40 |
| **Micronutrients** |  |  |
| Mn | MnCl<sub>2</sub>·4H<sub>2</sub>O | 0.5 |
| Mo | (NH<sub>4</sub>)<sub>2</sub>Mo<sub>7</sub>O<sub>24</sub>·4H<sub>2</sub>O | 0.05 |
| B | H<sub>3</sub>BO<sub>3</sub> | 0.2 |
| Zn | ZnSO<sub>4</sub>·7H<sub>2</sub>O | 0.01 |
| Cu | CuSO<sub>4</sub>·5H<sub>2</sub>O | 0.01 |
| Fe | FeCl<sub>3</sub>·6H<sub>2</sub>O | 2 |

### ^15^N stable isotope labeling

At 21 days after planting (DAP), differential ^15^N stable isotope labeling treatments were imposed for the final 24 h of plant growth to independently quantify acquisition of NH_4_^+^-N and NO_3_^-^-N. Ammonium and nitrate acquisition were traced using ^15^NH_4_NO_3_ and NH_4_^15^NO_3_, respectively (98 atom% ^15^N; Thermo Fisher Scientific). Labeled and unlabeled NH_4_NO_3_ were combined to provide a nominal 5 atom% ^15^N enrichment of the respective labelled N pool, with the quantity of labelled material adjusted to account for its 98 atom% ^15^N enrichment. The remaining NH_4_NO_3_ was supplied in unlabeled form. Thus, the two labeling solutions contained equivalent total N concentrations and NH_4_^+^:NO_3_^-^ ratios and differed only in whether the NH_4_^+^ or NO_3_^-^ pool was enriched with ^15^N. All other components of the Yoshida nutrient solution remained unchanged.

Fresh nutrient solution was prepared immediately prior to the labeling period. For each experimental run, 20 L of each labeling solution were prepared and divided equally between the two CO treatment chambers, providing 10 L per NFT tier. Existing nutrient solution was completely drained from each reservoir, and the reservoir was rinsed with deionized water prior to addition of the labeled solution. Within each growth chamber, one NFT tier received ^15^NH_4_NO_3_ - labeled solution and the other received NH_4_^15^NO_3_ - labelled solution. Assignment of isotopic labeling treatment to NFT tier was alternated between independent experimental runs to minimize potential confounding of labeling treatment with tier-specific effects.

Reservoir working volume was reduced from 20 to 10 L during the labeling period to reduce the quantity of ^15^N-labeled material required while maintaining continuous operation of the NFT system. Following addition of the labelled nutrient solution, pH was adjusted to 5.8 but was not subsequently corrected during the 24-h labeling period. All other environmental conditions were maintained as described above. Plants were exposed to the labeled nutrient solutions for exactly 24 h and harvested immediately following completion of the labeling period.

### Plant harvest and sample preparation

Plants were harvested at 22 days after planting (DAP), immediately following completion of the 24-h ^15^N labeling period. Shoots were excised at the surface of the foam cube and weighed immediately to determine shoot fresh weight. Roots were discarded because extensive root growth through the foam substrate prevented consistent recovery of intact root systems.

Individual shoots were placed into paper bags and oven-dried at 70 °C for 7 d. Following drying, shoot dry weight was recorded. The entire dried shoot from each individual plant was then transferred to a 50 mL centrifuge tube containing approximately 5 mL by volume of 5 mm glass beads and homogenized to a fine powder using a Bead Ruptor Elite bead mill homogenizer (OMNI International, Kennesaw, GA, USA). Samples were homogenized at 5 m s ¹ for two 30 s cycles separated by a 30 s dwell period to minimize sample heating.

Following homogenization, an aliquot of each powdered shoot sample was transferred to a 1.5 mL microcentrifuge tube.

### Isotope-ratio mass spectrometry and calculation of nitrogen metrics

Approximately 3 mg of homogenized shoot tissue from each plant was weighed into a tin capsule and analyzed for total N concentration and ^15^N enrichment at the University of Florida Stable Isotope Mass Spectrometry (SIMS) Laboratory. Samples were analyzed using a Costech ECS 4010 CHNS-O elemental analyzer coupled via a ConFlo IV interface to a Thermo Finnigan Delta V Advantage isotope-ratio mass spectrometer operating in continuous-flow mode. Samples underwent flash combustion at 1000 °C in an oxygen-rich atmosphere, followed by reduction at 650 °C and removal of water using a magnesium perchlorate trap. N_2_ and CO_2_ were subsequently separated using a 3.0 m gas chromatography column maintained at 55 °C prior to elemental and isotopic analysis. Tissue N concentration was determined as percentage N (%N) by comparison with calibration standards. For ^15^N-enriched samples, isotopic composition was reported as atom percent ^15^N (atom% ^15^N). A 1.0 atom% ^15^N ammonium sulfate standard and natural-abundance USGS40 reference material were included during analysis.

Differential ^15^N enrichment was used to assess relative acquisition of NH_4_^+^ and NO_3_^-^. Because plants supplied with ^15^NH_4_NO_3_ and NH ^15^NO_3_ received equivalent concentrations of NH ^+^ and NO ^-^ and differed only in the isotopically labelled N form, greater shoot atom% ^15^N following ^15^NH_4_^+^ labeling was interpreted as greater acquisition of ammonium-derived N, whereas greater enrichment following ^15^NO_3_^-^ labeling indicated greater acquisition of nitrate-derived N.

Total shoot N content was calculated for each individual plant from shoot dry mass and tissue N concentration determined by elemental analysis as:

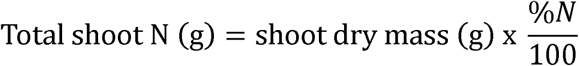

### Statistical analysis

Statistical analyses were performed in R using linear mixed-effects models fitted with the lmerTest package. Individual plants represented biological observations; however, plants within a common NFT tier shared the same recirculating nutrient reservoir and were therefore treated as subsamples of a common experimental unit. NFT tier, uniquely defined within experimental run and growth chamber, was included as a random intercept to account for this hierarchical structure. Experimental run was included as a fixed blocking factor because the experiment was conducted in three independent runs over time. Type III tests of fixed effects were performed using Satterthwaite approximations for denominator degrees of freedom. Model assumptions were assessed by visual inspection of residual-versus-fitted and normal Q–Q plots. Statistical significance was assessed at α = 0.05.

Shoot ^15^N enrichment (atom% ^15^N) was analyzed separately for each lettuce cultivar using models containing isotopic label (^15^NH ^+^ or ^15^NO ^-^), CO concentration (410 or 3000 ppm), and their interaction as fixed effects, with experimental run included as a fixed block and NFT tier as a random intercept. Estimated marginal means were calculated using the emmeans package. Planned pairwise comparisons were performed between isotopic labels within each CO_2_ concentration and between CO_2_ concentrations within each isotopic label, with *P*-values adjusted using the Holm method. Model-estimated marginal means and 95% confidence intervals were used to summarize treatment responses. To visualize the magnitude and direction of N-form preference, a preference index was calculated from the model-estimated marginal means as ^15^N_NO3_ − ^15^N_NH4_, with negative values indicating greater acquisition of NH ^+^-derived N and positive values indicating greater acquisition of NO ^-^-derived N. Standard errors for the preference index were obtained from the corresponding model-estimated contrast. Statistical inference concerning N-form acquisition was based on the underlying mixed-effects models and planned contrasts rather than significance testing of the preference index as a separate response variable.

Shoot fresh weight and total shoot N content were analyzed using linear mixed-effects models containing lettuce cultivar, CO_2_ concentration, and their interaction as fixed effects. Isotopic label and experimental run were included as additive fixed effects, and NFT tier was included as a random intercept. Isotopic label was retained in these models to account for assignment of the isotopic treatments, although the two labeling solutions were chemically equivalent apart from the isotopically enriched N form. Where appropriate, estimated marginal means were compared using Tukey-adjusted pairwise comparisons. For total shoot N, additional planned comparisons between CO_2_ concentrations within each cultivar were performed using Holm adjustment. Figures show individual plant observations and/or treatment distributions together with model-derived summaries or multiple-comparison groupings, as appropriate.

## Results

### Differential ^15^N labeling reveals consistent ammonium preference across lettuce cultivars

Shoot ^15^N enrichment differed significantly between isotopic labeling treatments in all four lettuce cultivars, indicating differential acquisition of the two inorganic N forms. The effect of isotopic label was significant for Dragoon, Outredgeous, Waldmann’s Dark Green, and Black Seeded Simpson. By contrast, neither CO_2_ concentration, nor the Label × CO_2_ interaction were significant for any cultivar (Fig. 1).

**Figure 1:**
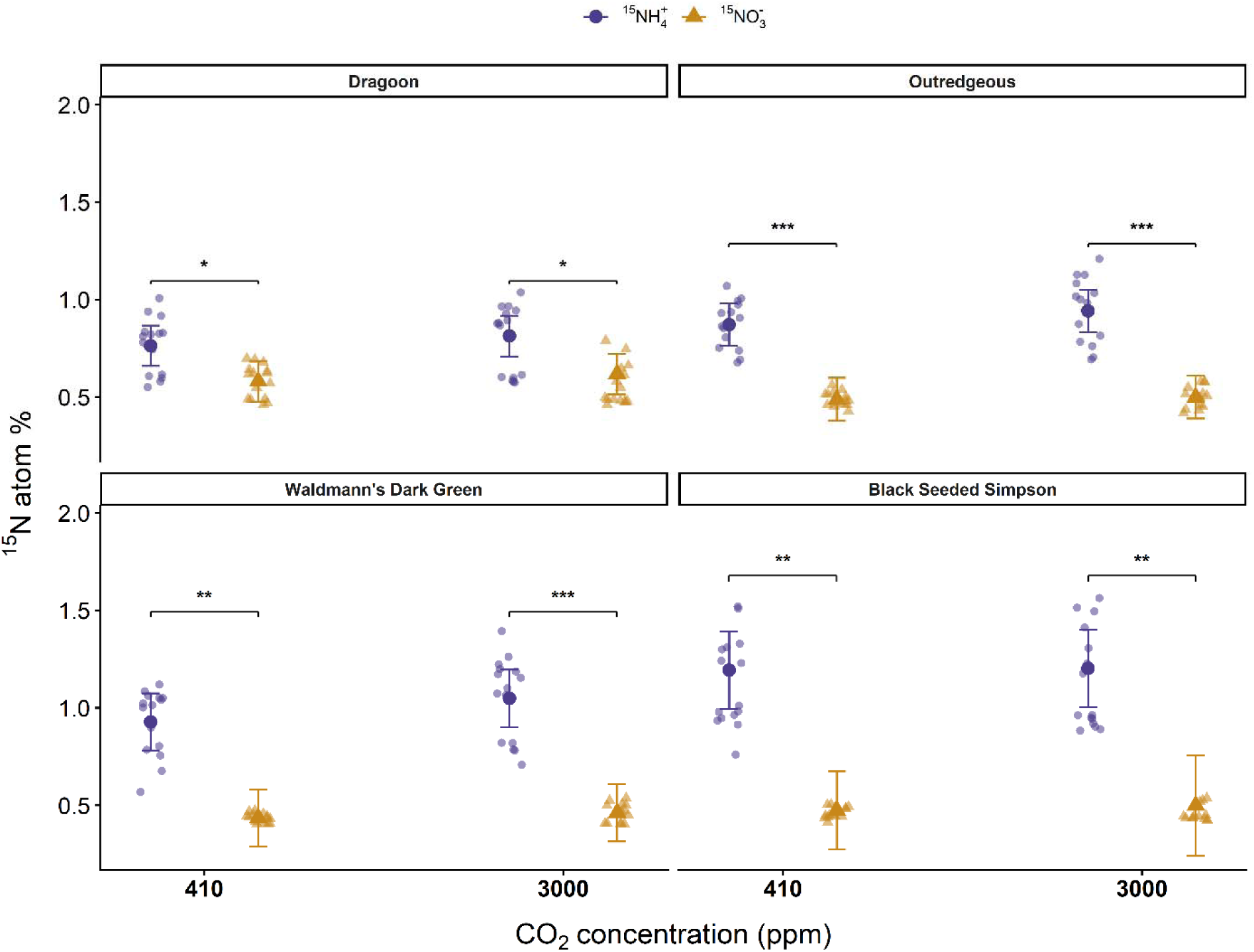
Differential acquisition of ammonium- and nitrate-derived nitrogen by lettuce under contrasting CO_2_ environments. Shoot ^15^N enrichment of four lettuce (*Lactuca sativa* L.) cultivars following differential ^15^NH_4_^+^ or ^15^NO_3_^-^ labeling under 410 and 3000 ppm CO_2_. Plants were exposed to isotopically labelled nutrient solutions for the final 24 h of growth. Small points represent individual plants and larger points with error bars represent estimated marginal means ± 95% confidence intervals derived from linear mixed-effects models fitted separately for each cultivar. Models included isotopic label, CO_2_ concentration, and their interaction as fixed effects, experimental run as a fixed block, and NFT tier as a random intercept. Solid brackets indicate significant pairwise differences between ^15^NH_4_^+^- and ^15^NO_3_^-^ - labeled treatments within a CO_2_ concentration following Holm adjustment. Only significant comparisons are shown. P < 0.05 (*), P < 0.01 (**), and P < 0.001 (***).

Within each CO_2_ environment, shoot ^15^N enrichment was consistently greater following ^15^NH_4_^+^ than ^15^NO_3_^-^ labeling. At 410 ppm CO_2_, the model-estimated ^15^NH_4_^+^ − ^15^NO_3_^-^ differences were 0.182 atom% for Dragoon, 0.383 atom% for Outredgeous, 0.493 atom% for Waldmann’s Dark Green, and 0.718 atom% for Black Seeded Simpson. The same pattern was retained at 3000 ppm CO_2_, with corresponding differences of 0.195, 0.443, 0.587, and 0.703 atom%, respectively in all cases (Fig. 1). No within-label comparison between 410 and 3000 ppm CO_2_ was significant after Holm adjustment.

The model-derived N-form preference index was negative for all cultivars under both CO_2_ conditions, confirming greater acquisition of NH_4_^+^-derived than NO_3_^-^-derived N (Fig. 2). The magnitude of ammonium preference increased in the order Dragoon < Outredgeous < Waldmann’s Dark Green < Black Seeded Simpson. Preference indices at 410 and 3000 ppm CO_2_, respectively, were −0.182 and −0.195 for Dragoon, −0.383 and −0.443 for Outredgeous, −0.493 and −0.587 for Waldmann’s Dark Green, and −0.718 and −0.703 for Black Seeded Simpson. Although Dragoon, Outredgeous and Waldmann’s Dark Green showed numerically greater ammonium preference at 3000 ppm, the absence of significant Label × CO_2_ interactions indicated that CO_2_ concentration did not significantly alter N-form preference.

**Figure 2:**
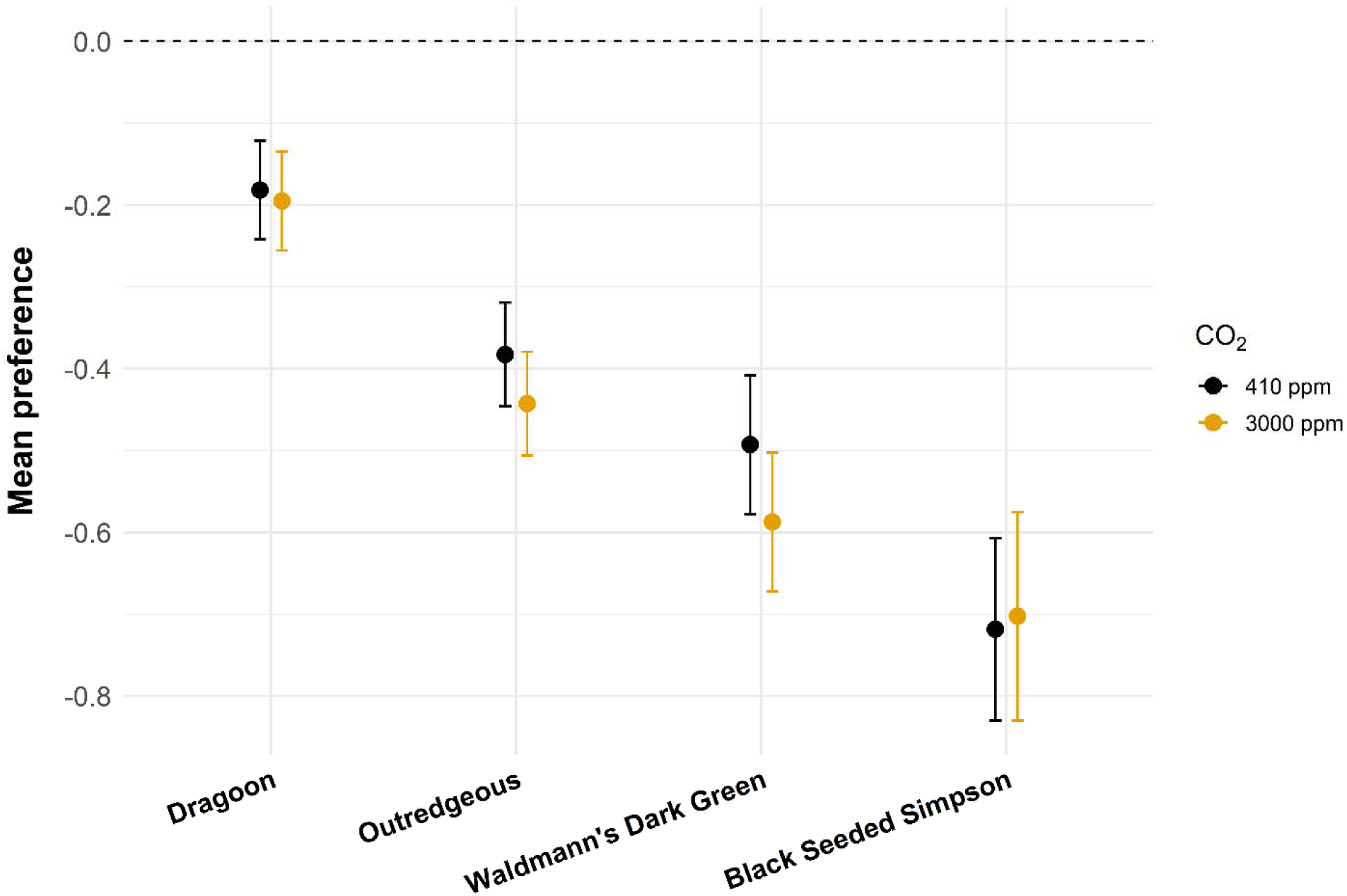
Nitrogen-form preference differs among lettuce cultivars. Nitrogen-form preference of four lettuce (*Lactuca sativa* L.) cultivars grown under 410 and 3000 ppm CO_2_. Preference indices were calculated from model-estimated marginal means as shoot ^15^N enrichment following ^15^NO_3_^-^ labeling minus enrichment following ^15^NH_4_^+^ labeling. Negative values indicate greater acquisition of NH_4_^+^-derived N, positive values indicate greater acquisition of NO_3_^-^-derived N, and the dashed horizontal line at zero represents equivalent acquisition of the two N forms. Points represent model-estimated preference indices and error bars represent standard errors of the corresponding model-estimated contrasts. The preference index is presented as a summary visualization; statistical inference concerning N-form acquisition was performed on the underlying ^15^N enrichment data (Fig. 1).

### Shoot fresh biomass differs among cultivars but not CO_2_ environments

Shoot fresh weight differed significantly among lettuce cultivars, whereas there were no significant effects of CO_2_ concentration, isotopic label, or Cultivar × CO_2_ interaction (Fig. 3). Waldmann’s Dark Green and Black Seeded Simpson produced the greatest shoot biomass, with estimated marginal means of 13.91 and 12.87 g plant ¹, respectively, and did not differ significantly from one another. Both cultivars had significantly greater shoot fresh biomass than Dragoon (10.14 g plant ¹) and Outredgeous (9.02 g plant ¹), which did not differ significantly from one another.

**Figure 3.**
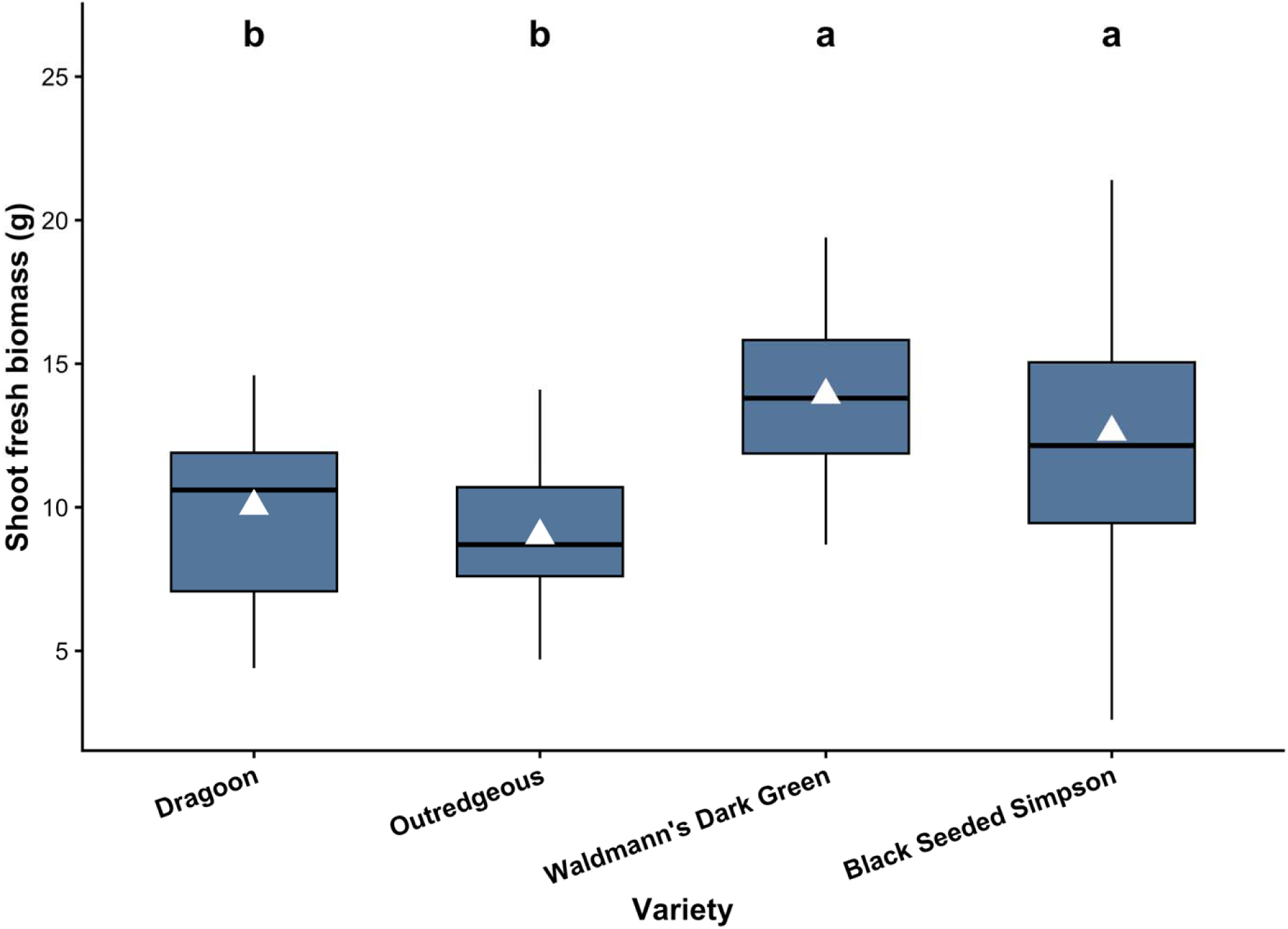
Shoot fresh biomass differs among lettuce cultivars. Shoot fresh weight of four lettuce (*Lactuca sativa* L.) cultivars following 22 d of hydroponic growth. Boxplots show the distribution of individual plant observations pooled across CO_2_ an isotopic labeling treatments; horizontal lines indicate medians, boxes represent the interquartile range, and whiskers extend to 1.5 × the interquartile range. Triangles indicate arithmetic means. Data were analyzed using a linear mixed-effects model containin cultivar, CO_2_ concentration, and their interaction as fixed effects, isotopic label and experimental run as additive fixed effects, and NFT tier as a random intercept. Different letters indicate significant differences among cultivar estimated marginal means following Tukey adjustment (*P* < 0.05).

### Total shoot nitrogen content shows a cultivar-dependent response to CO_2_ environment

Total shoot N content differed significantly among cultivars and showed a significant Cultivar × CO_2_ interaction (Fig. 4). Neither the overall effect of CO_2_ concentration, nor isotopic label were significant.

**Figure 4.**
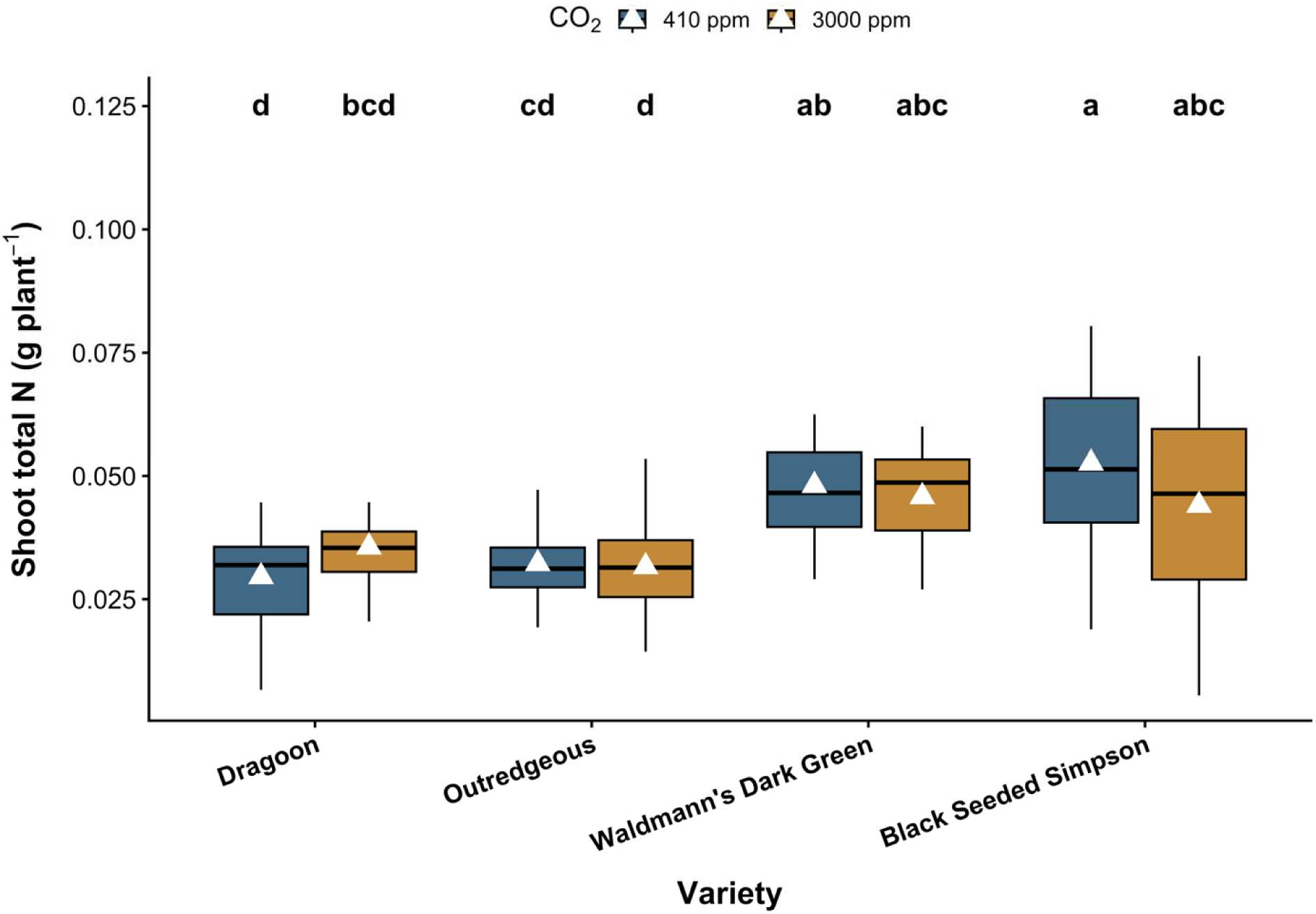
Total shoot nitrogen content varies with lettuce cultivar and CO□ environment. Total shoot N content of four lettuce (*Lactuca sativa* L.) cultivars grown under 410 or 3000 ppm CO_2_. Total shoot N was calculated from shoot dry mass and tissue N concentration determined by elemental analysis. Boxplots show individual plant distributions; horizontal lines indicate medians, boxes represent the interquartile range, and whiskers extend to 1.5 × the interquartile range. Triangles indicate arithmetic means. Data were analyzed using a linear mixed-effects model containing cultivar, CO_2_ concentration, and their interaction as fixed effects, isotopic label and experimental run as additive fixed effects, and NFT tier as a random intercept. Different letters indicate significant differences among cultivar × CO_2_ estimated marginal means following Tukey adjustment (*P* < 0.05).

The highest model-estimated total shoot N content occurred in Black Seeded Simpson at 410 ppm CO_2_ (0.0526 g plant^-1^), followed by Waldmann’s Dark Green at 410 ppm (0.0481 g plant^-1^) and 3000 ppm (0.0458 g plant^-1^) and Black Seeded Simpson at 3000 ppm (0.0445 g plant^-1^). Dragoon and Outredgeous generally accumulated less total shoot N. Despite the significant overall Cultivar × CO_2_ interaction, Holm-adjusted comparisons between CO_2_ concentration within individual cultivars were not significant. Black Seeded Simpson showed the largest numerical difference between CO_2_ environments, with total shoot N 0.0081 g plant^-1^ greater at 410 than 3000 ppm, whereas the corresponding within-cultivar differences were smaller for Dragoon, Outredgeous, and Waldmann’s Dark Green.

## Discussion

The present study demonstrates a consistent preference for ammonium-derived N across four lettuce cultivars when NH_4_^+^ and NO_3_^-^ were supplied simultaneously at equivalent concentrations.

Shoot ^15^N enrichment was significantly greater following ^15^NH_4_^+^ than ^15^NO_3_^-^ labeling in every cultivar and under both CO_2_ environments. This contrasts with the nitrate-dominant nutrition conventionally employed in hydroponic crop production (Sonneveld & Voogt, 2009; Voogt & Bar-Yosef, 2019) but is physiologically plausible. NO_3_^-^ must be reduced to NO_2_^-^ and subsequently NH_4_^+^ before entering the GS/GOGAT assimilation pathway, whereas absorbed NH_4_^+^ can be assimilated directly, avoiding the additional energetic and reductant requirements associated with NO_3_^-^ reduction (Bloom et al., 1992). Although NH_4_^+^ nutrition carries additional physiological costs associated with ion homeostasis, carbon-skeleton provision and pH regulation (Britto & Kronzucker, 2013), these results demonstrate preferential NH acquisition when both inorganic N forms are similarly supplied.

This distinction between physiological acquisition and environmental availability is central to interpreting N-form preference. Studies in natural systems demonstrate that apparent preference can shift with environmentally mediated N availability, both across communities and within species (Houlton et al., 2007; Wang & Macko, 2011), while the concept of fixed N-form preference has been questioned because differential acquisition in soil can reflect N transformations and differences in availability rather than plant discrimination alone (Chalk & Smith, 2021). Hydroponics controls many of these environmental determinants, allowing NH_4_^+^ and NO_3_^-^ to be supplied directly and at defined concentrations. Our differential-labeling approach further minimized differences in external supply, yet all four cultivars preferentially acquired NH_4_^+^. Consequently, the widespread use of NO_3_^-^ as the dominant N source in hydroponic crop production should not necessarily be interpreted as evidence that it represents the intrinsically preferred N form of lettuce.

Elevated CO_2_ was hypothesized to strengthen NH_4_^+^ preference because elevated CO_2_ can inhibit shoot NO_3_^-^ assimilation in C_3_ plants and alter responses to N form (Bloom et al., 2002; Bloom et al., 2012). Contrary to this hypothesis, CO_2_ concentration did not significantly alter N-form preference. One possibility is that the strong NH_4_^+^ preference already present at 410 ppm provided limited scope for a further shift. Alternatively, changes in shoot NO_3_^-^ assimilation may not translate directly into whole-plant acquisition when both N forms remain continuously available. Importantly, NH_4_^+^ preference persisted in all cultivars at 3000 ppm, a spaceflight-relevant elevated CO_2_ concentration within the range historically encountered aboard the ISS (Beard, 2020; Georgescu et al., 2020). The CO_2_ comparison should nevertheless be interpreted cautiously because each concentration was maintained in a dedicated growth chamber, preventing separation of CO_2_ from persistent chamber effects. The strongest inference therefore lies in the replicated differential-label comparisons within each environment: lettuce preferentially acquired NH_4_^+^ under both ambient and highly CO_2_-enriched conditions.

The magnitude of NH_4_^+^ preference varied substantially among cultivars and broadly corresponded with biomass and total shoot N accumulation. Dragoon exhibited the weakest preference and comparatively low biomass and shoot N, whereas Black Seeded Simpson exhibited the strongest preference alongside greater biomass and N accumulation. Importantly, this gradient was driven predominantly by increasing ^15^N enrichment following ^15^NH_4_^+^ labeling rather than decreasing ^15^NO_3_^-^ enrichment. A simple effect of plant size would be expected to influence acquisition from both labelled pools, while atom% ^15^N represents proportional isotopic composition rather than total tracer mass. The data therefore are consistent with a comparatively stable contribution from NO_3_^-^ acquisition accompanied by increasing incorporation of NH_4_^+^-derived N as plant growth and N demand increase. Shoot atom% ^15^N does not provide an absolute uptake flux, so this mechanism cannot be demonstrated conclusively; however, it suggests a testable hypothesis that rapidly growing or highly N-demanding genotypes may increasingly satisfy additional N demand through NH_4_^+^ acquisition when both forms are available.

These findings have implications for nutrient management in space crop production, where crop performance must ultimately be considered alongside resource use and integration with regenerative life-support systems. Nutrient-loop closure and recycled fertigation have been identified as important steps toward operational BLSS (Fountain et al., 2026). Increasing the proportion of N supplied as NH_4_^+^ could better align nutrient provision with lettuce acquisition physiology, although whether this improves yield or N-use efficiency requires direct testing across optimized NH_4_^+^:NO_3_^-^ ratios. Recent work showing improved hydroponic lettuce performance when NH comprised 20% of supplied N further supports optimization of crop-specific N-form ratios rather than assuming nitrate-dominant nutrition is optimal (Xu et al., 2025).

N-form preference may become particularly important when crop production is coupled with nutrient recovery. Urinary N is predominantly excreted as urea and converted to NH_4_^+^ following hydrolysis, whereas producing NO_3_^-^ from this recovered N requires nitrification (Verbeelen et al., 2021). Greater direct utilization of recovered NH_4_^+^ could therefore potentially reduce the extent of processing required before crop delivery and help integrate crop production with nutrient-loop closure (Verbeelen et al., 2021; Fountain et al., 2026). Similar opportunities exist in terrestrial CEA, where ammonium-rich recovered nutrient streams such as anaerobic-digestion digestate could be integrated with hydroponic crop production (van Rooyen & Nicol, 2022). Preferential NH_4_^+^ acquisition should not, however, be interpreted as evidence that unrestricted or sole-source NH_4_^+^ nutrition is optimal. Excessive NH can impair growth and drive rapid root-zone acidification (Britto & Kronzucker, 2013; Natarajan Sakthivel et al., 2023), making optimization of NH_4_^+^:NO_3_^-^ supply and pH management essential.

More broadly, these results demonstrate that nitrate-dominant nutrition does not necessarily reflect intrinsic N-form acquisition preference. Engineered hydroponic systems provide an opportunity to approach N nutrition differently from soil-based agriculture: rather than allowing environmental processes to determine the N forms available to roots, nutrient supply can be designed around crop acquisition physiology. The consistent NH_4_^+^ preference observed across four lettuce cultivars provides a physiological basis for reconsidering N-form provision in CEA and for integrating crop nutrient requirements with locally available and recovered N resources on Earth and beyond.

## Acknowledgements

L.L.F. was supported by an appointment to the NASA Postdoctoral Program at NASA Kennedy Space Center, administered by Oak Ridge Associated Universities under contract with NASA. The authors thank the Kennedy Space Center LASSO-II team for assistance with experimental activities and the University of Florida Stable Isotope Mass Spectrometry Laboratory for isotope-ratio mass spectrometry analyses.

## Author Contributions

L.L.F., E.B.B., R.M.W., and G.D.M. conceived and designed the experiments. L.L.F. performed the experiments. L.L.F. and S.M. analyzed the data. L.L.F. drafted the manuscript with contributions from S.M. E.B.B., R.M.W., and G.D.M. critically reviewed and revised the manuscript. All authors approved the final manuscript.

